# Vimentin provides the mechanical resilience required for amoeboid migration and protection of the nucleus

**DOI:** 10.1101/720946

**Authors:** Luiza Da Cunha Stankevicins, Marta Urbanska, Daniel AD. Flormann, Emmanuel Terriac, Zahra Mostajeran, Annica K.B. Gad, Fang Cheng, John E. Eriksson, Franziska Lautenschläger

## Abstract

Dendritic cells use amoeboid migration through constricted passages to reach the lymph nodes, and this homing function is crucial for immune responses. Amoeboid migration requires mechanical resilience, however, the underlying molecular mechanisms for this type of migration remain unknown. Because vimentin intermediate filaments (IFs) and microfilaments regulate adhesion-dependent migration in a bidirectional manner, we analyzed if they exert a similar control on amoeboid migration. Vimentin was required for cellular resilience, via a joint interaction between vimentin IFs and F-actin. Reduced actin mobility in the cell cortex of vimentin-reduced cells indicated that vimentin promotes Factin subunit exchange and dynamics. These mechano-dynamic alterations in vimentin-deficient dendritic cells impaired amoeboid migration in confined environments *in vitro* and blocked lymph node homing in mouse experiments *in vivo*. Correct nuclear positioning is important in confined amoeboid migration both to minimize resistance and to avoid DNA damage. Vimentin-deficiency also led to DNA double strand breaks in the compressed dendritic cells, pointing to a role of vimentin in nuclear positioning. Together, these observations show that vimentin IF-microfilament interactions provide both the specific mechano-dynamics required for dendritic cell migration and the protection the genome needs in compressed spaces.

**Summary statement:** Vimentin — in joint action with actin — mediates the mechanical stiffness of cells required for amoeboid cell migration through confined spaces and protects the nucleus from DNA damage.

## Introduction

Vimentin is a member of the large intermediate filament (IF) protein family, which is characterized by a remarkable diversity in terms of protein sequences, expression patterns, and the distribution in various tissues [1]. Vimentin is the major IF protein in a broad variety of cell types, especially in motile and dynamic cells of mesenchymal origin. In this regard, vimentin has been implicated to be involved in different migratory functions of a number of different adherent cell types, especially in relation to the organization and functionalities of actomyosin complexes [2–4]. The role of vimentin in cell migration has been primarily addressed in adherent cells [5]. However, the role of vimentin in the migration of low-adherent or suspended cells remains unknown.

Among all low-adherent cells in the body, the immune cells are of particular importance. Cells of the immune system migrate in an amoeboid mode of migration, which is fundamentally different to the more commonly studied mesenchymal mode of migration, which is characterized by actin stress fibers and focal adhesions to the extracellular matrix [6]. In comparison, amoeboid migration is a rapid mode of motility that is characterized by minimal adhesion and high contractility [6]. Ameboid migration depends on friction forces with the environment that might be compared to those employed by persons engaged in chimney climbing [7].

Mice that lack vimentin have immune defects [8]. These defects have been suggested to be caused by defective lymphocyte migration [9], however, the role of vimentin in the molecular control of amoeboid migration of the involved cell types remains unclear.

The mechanical behavior of a migrating immune cell is mainly determined by the cytoskeleton, a complex and dynamic structure based on filamentous actin, microtubules, and IFs, including also cytoskeletal crosslinkers and molecular motor proteins. In addition, the nucleus also provides mechanical support for migrating immune cells [10]. The roles of the microfilaments and microtubules, and their control of the mechanical properties of cells have been extensively studied (for overview, see [10]). In contrast, the role of IFs in the control of the mechanical properties of cells has only been recently investigated [1, 11] and details of the mechanism behind the involvement of vimentin are still lacking. Most of these studies have relied on cells that adhered to a 2D surfaces, and even less studies address the role of vimentin IFs in low-adherent cells. We have previously shown that the mechanical properties of suspended and adherent cells are fundamentally different [12]. In suspended cells, the cytoskeleton is adapted to the low-adhesive state of the cells, and the cytoskeletal control of cellular mechanics is likely to be different in suspended than in adherent cells. The collapsed, perinuclear localization of vimentin in lymphocytes suggests that vimentin may provide mechanical stability to the cell body enabling transmigration through the endothelium [13]. Therefore, we hypothesized that the vimentin network governs the mechanical stiffness and the forces that underly the amoeboid migration of low-adherent immune cells.

In this study, we aim to clarify if the vimentin network can control the dynamic cell stiffness properties required for immune cells to be able to efficiently migrate through confined spaces. As a model system, we chose bone marrow-derived dendritic cells (BMDC), because of their capacity to migrate through narrow passages is required for the innate immune responses. We analyzed BMDC migration both *in vitro* and *in vivo* by measuring homing to lymph nodes. Our results show that vimentin IFs together with microfilaments provide the biomechanical properties required for dendritic cell migration *in vitro* and *in vivo*, and suggest a role for vimentin in protecting the genome integrity during the nuclear compression of migrating dendritic cells.

## Results

Adaptive immune responses rely on a rapid, spatiotemporally concerted migration of BMDCs to the lymph nodes or to the site of inflammation [14]. Although many parts of these process are well understood, little is known about the detailed molecular mechanisms underlying the actual migration and passage through complex tissue settings and highly confined spaces. It is clear that the involved migratory processes require active engagement of the cytoskeleton, as a number of studies have demonstrated specific roles of microtubules and microfilaments. However, less is known about the functions of IFs, although vimentin has been demonstrated to play an essential role in leukocyte homing [5]. Therefore, we wished to specifically determine the contribution of vimentin to the biomechanics required for BMDC migration and homing.

### Vimentin is required for the normal rate of migrating BMDC

The confinement of cells has been shown to be crucial for the migration properties of BMDCs [15]. Therefore, we wanted to analyze the migration of these cells *in vitro*, using conditions that mimic the confined, *in vivo* physiological environment of BMDCs. To this end, we produced 1D confining channels (Fig. 1A, B) and 2D confining plates (Fig. 1C, D), as described in detail in the Materials and Methods section and in previous publications [16–18]. These devices were then used to analyze the capacity of cells to migrate in 1D and 2D. Overall, we observed a marked reduction of cell migration upon loss of vimentin (Fig. 1G-K, table S1). This reduction was observed in terms of a decreased total number of migrating cells (Fig. 1G) as well as in a decreased migration speed (Fig. 1H), and was observed both in 1D and 2D migration. In addition, we observed a reduced lengths of migration paths in vimentin-deficient cells (Fig 1I). However, loss of vimentin did not result in any significant changes of the apparent persistence of cells in our system (Fig. 1J). Taken together, these data indicate that vimentin is essential for the BMDCs to migrate efficiently in a 1D or 2D confined environment.

**Figure 1.**
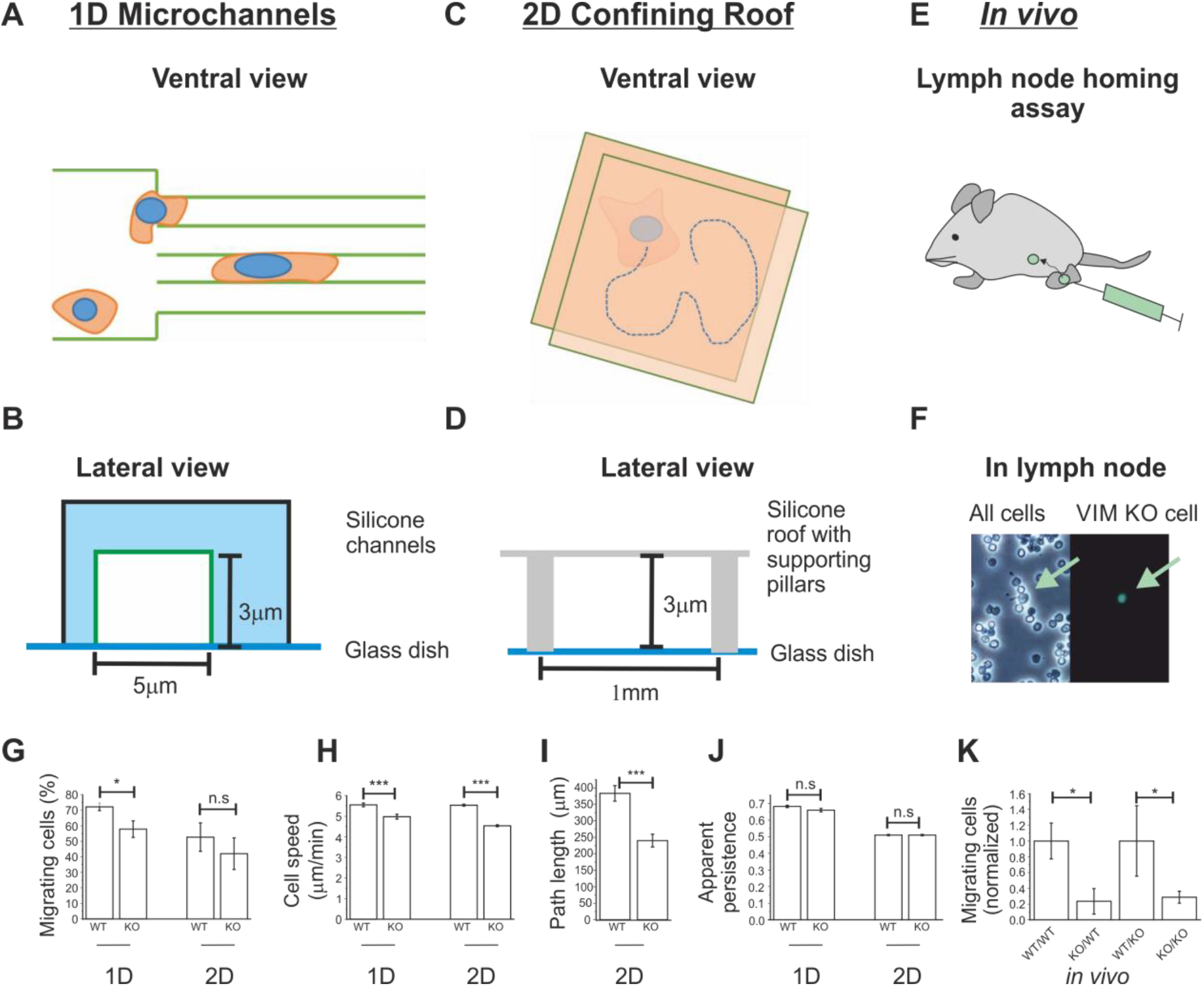
Loss of vimentin results in defective amoeboid migration. Ventral (A) and lateral (B) view of scheme of 1D channel migration set-up, ventral (C) and lateral (D) view of scheme of 2D roof set-up on. (E) Scheme of lymph node homing assay. (F) Representative phase contrast (left) and fluorescence (right) microscopy images, respectively, used to count for fluorescently labeled VIM KO cells arrived at the lymph node. Green arrow points at one VIM KO cell taken from an experiment when VIM KO cells were injected. (G) percentage of migrating BMDC wt and BMDC vimentin KO in 1D and 2D, (H) speed of migrating BMDC WT and BMDC vimentin KO in 1D and 2D. (I) Path length of migration trajectories of migrating BMDC wt and BMDC vimentin KO 2D (1D not applicable since given by the length of the channel) (J) Apparent persistence of migrating BMDC WT and BMDC vimentin KO in 1D and 2D. (K) Normalized ratio of injected BMDC WT and BMDC vimentin KO arriving at lymph node either in wt (wt/wt and ko/wt) or in vimentin KO mice (wt/ko and ko/ko). * *p* ≤0,05. Bars represent mean value SE.

### Vimentin is required for the migration of dendritic cells to the lymph node

In order to validate the *in vitro* results above, we wished to analyze if vimentin controls BMDCs migration in their physiological context when dendritic cells migrate to lymph nodes after an encounter with pathogens. To this end, we analyzed the capacity of LPS-treated dendritic cells to migrate to the lymph nodes using an *in vivo* lymph node-homing assay, as described in the Materials and Methods section and in previous publications [19]. We injected LPA-treated dendritic cells in the footpad of mice and analyzed the amount of BMDCs cells in the closest lymph node after 36 hours (Fig. 1F). We observed that the homing of vimentin-deficient BMDCs in wt mice was significantly impaired as compared to wt BMDCs (Fig. 1K). Identical results were obtained in vimentin-deficient mice. These data indicate that the observed deficiency is due to the homing capacity of vimentin-deficient BDMCs and not due to the vimentin-levels in the surrounding microenvironment and/or receiving tissue(s). Taken together, these observations support the results we obtained in our *in vitro* models, and demonstrate that vimentin is essential for BMDCs to reach the lymph nodes.

### Vimentin is important for the elastic but not the viscous properties of BMDCs cells

We have previously shown that cell mechanics and cell migration are closely interdependent phenomena [20], and vimentin has been reported to control the mechanical properties of cells ([1, 5]). When determining the subcellular localization of actin and vimentin IFs in immature BMDCs 24h after seeding on glass, we observed a predominantly perinuclear subcellular localization of vimentin filaments, with no detectable vimentin at the cell periphery or in cell protrusions (Fig. S1). During migration through tissues, the cell body of dendritic cells are confined between other cells and extracellular matrix [15, 21]. To determine the subcellular localization of vimentin in a context that better mimics the *in vivo*-situation, we imaged cells in custom-engineered 1D confining channels [15, 22]. We observed that also here, vimentin IFs were predominantly localized close to the nucleus, with no signal at the periphery or in cellular protrusions (Fig. SI 1). These observations are in line with earlier reports that vimentin IFs localize predominantly around the nucleus in cells (see [1] for overview), whereas peripheral cell parts show less prominent IF structures.

Because the BMDCs showed a predominantly perinuclear localization of vimentin IFs (Fig. S 1), we hypothesized that vimentin mainly controls the mechanical properties of the central, perinuclear region, and not of the peripheral cortex of these cells. To test this hypothesis, we analyzed the mechanical properties at both subcellular locations of BMDCs with or without expression of vimentin. To measure whole-cell mechanical properties under low deformations, we implemented a high-throughput method that deforms cells with a hydrodynamic shear flow on a millisecond timescale, called real-time deformability cytometry (RT-DC) ([23], Fig. 2A), and therefore probes predominantly the outer, cortical cell region. The applied shear flow can be adapted to tune the deformation forces imposed on the cells. We used two flow rates*flow rate 1* (Fr1 = 0.16 μl s^-1^), causing low deformation of cells, and two-fold higher *flow rate 2* (Fr2 = 0.32 μl s^-1^), causing a higher degree of cell deformation. We observed only a minor difference in the deformation values between vimentin-deficient and control cells (Fig. 2B – D, table S2). This difference was statistically significant only for *flow rate 2*. Since in a microfluidic channel of fixed dimension the amount of force experienced by cells of dissimilar sizes is different, it is necessary to extract a material property, such as the Young’s modulus, in order to draw a comparative conclusion about mechanical properties of the cells measured, The Young’s modulus describes elastic properties of the cells and expresses how much stress is required to deform a cell by a certain amount A material with a high Young’s modulus is often described as *stiff*, and with a low Young’s modulus as *soft*. In order to calculate the Young’s modulus of the BMDCs deformed using RT-DC, we also measured the cell cross section and observed that the vimentin-deficient cells are significantly smaller (Fig. 2F). Taking into account the smaller area of the cells (Fig. 2F), the calculated Young’s modulus was significantly lower in vimentin-deficient cells compared to control cells for both flow rates applied (Fig. 2C, E).

**Figure 2.**
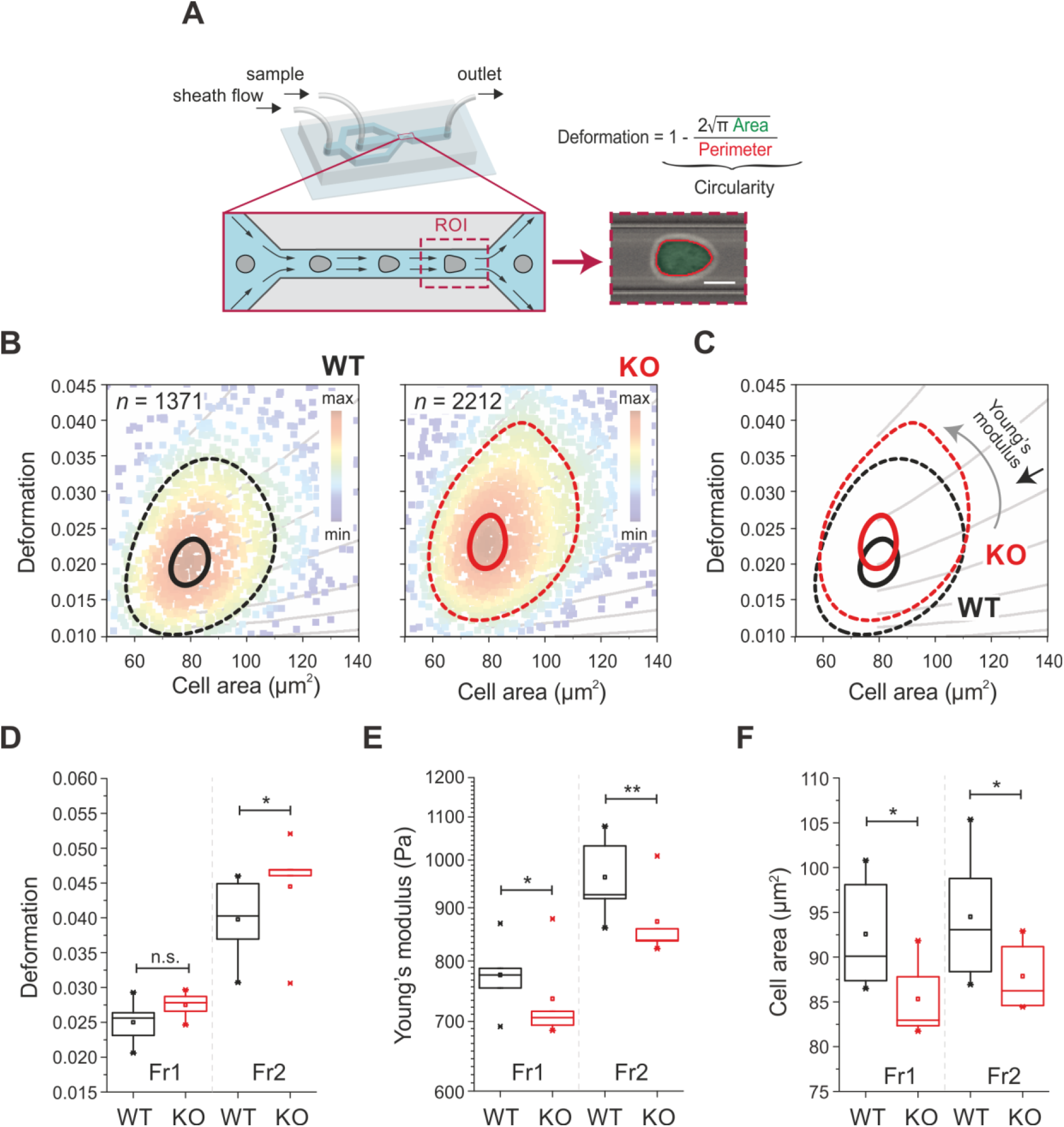
Loss of vimentin in dendritic cells decreases cell stiffness as detected by RT-DC. (A) Deformability measurements with RT -DC are performed in a microfluidic chip (shown in the background) at the end of a channel constriction (zoom -in). Cell deformation is evaluated using image-derived parameters. (B) Deformation -cell area scatter plots showing a representative measurement of wild type (wt) and vimentin knock -out (KO) cells. Cell number, *n*, is indicated on the corresponding plots. Color map indicates event density. Contour plots delineate 50% density (dashed lines) and 95% density (solid lines). (C) An overlay of contours from B. In B -C, grey lines delineate isoelastic regions obtained with the numerical simulations grouping cells of same mechanical properties. (D-F) Comparison of deformation (D), Young’s Modulus (E) and cell area (F) of wt and KO cells measured at two different flowrates: Fr1 = 0.16 μl s-1, and Fr2 = 0.32 μl s-1. In D -F, box plots of medians from 5 independent experiment replicates are shown, boxes present 25th to 75th percentile range, with a line at the median. Whiskers indicate extreme data points within 1.5 × the interquartile range (IQR). \*\**p* < 0.01; \**p* < 0.05; ns, not significant. Statistical analysis was performed using a linear mixed effects model.

To clarify the role of vimentin in the determination of the mechanical properties of the perinuclear, cytoplasmic region of BMDC cells, we used a low throughput method which offers better control of speed and depth of indentation compared to RT-DC: Atomic Force Microscopy (AFM). To this end, cells were placed on a non-adhesive PEG-coated dish, followed by a global measurement of cell mechanical properties, using wedged tiples cantilevers (Fig. 3A, B), as described in Materials and Methods section [24]. We measured cell force relaxations while compressing the cell successively five times, while keeping the indentation of the cantilever constant for 60 s per step and then lowered 1 μm on each step (Fig. 3 C). This analysis of the deformation curves, as described in Materials and Methods section, allows to evaluate the Young’s modulus to describe elastic parameters but also the relaxation time in order to estimate the viscosity. A higher relaxation time would indicate a higher viscosity. AFM is, therefore, ideally suited to describe the viscoelastic properties of cells. The first of the five indentations showed a lower Young’s modulus in vimentin-deficient cells as compared to control cells (Fig. 3D, table S3), indicating a lower elasticity in the vimentin-deficient cells. The deeper indentations showed the same reduced Young’s modulus in vimentin-deficient cells (Fig. 3D). In contrast, vimentin-deficient cells showed no difference in their relaxation time, as compared to control (Fig. 3F, table S3). Hence, as there was a difference in elasticity but not in viscosity, these data suggest that vimentin is important for the elastic, but not the viscous properties of suspended BMDCs.

**Figure 3.**
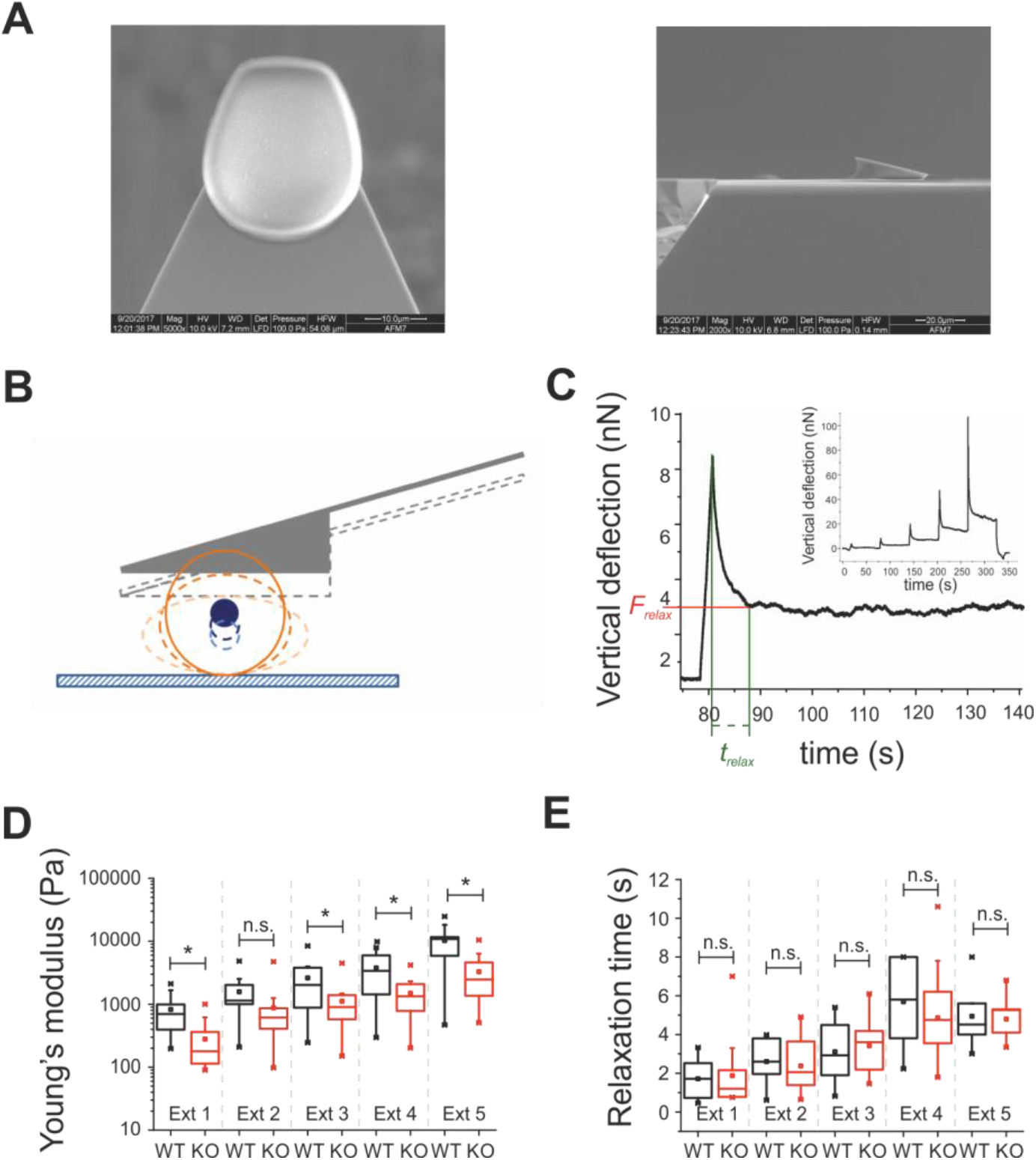
Force-mode atomic force microscopy analysis of dendritic cell mechanics. (A) Ventral and lateral electron microscope images of wedged cantilevers used to evaluate the cell global mechanical response by AFM. (B) Scheme of wedged cantilever compressing cell. (C) Scheme of deformation graph used for analysis, describing how relaxation time was measured. (D) Young’s modulus of BMDC wt (black) and vimentin KO (red) measured at different extends. (E) Relaxation time of BMDC wt and vimentin KO measured at different extends: Results of 3 independent experiments (total of 12 wt cells and 16 vim KO cells). * *p* ≤0,05. Whisker middle line represent median values. In boxplot mean values are represented by a square.

### Filamentous actin controls the modest and rapid but not the major and global mechanical response both in normal and vimentin-deficient BMDCs

Because actin microfilaments are known to be a prime determinant for mechanical properties in cells, we aimed to clarify the role of filamentous actin on the mechanical properties of the suspended BMDCs. To this end, we depolymerized actin in vimentin-deficient and control BMDCs, and analyzed the mechanical properties using both RT-DC and AFM. The analysis using RT-DC, measuring primarily the cortical stiffness, showed that upon loss of F-actin, the cells became significantly softer, and that this decreased stiffness was independent of vimentin for both flow rates applied (Fig. 4A-C, table S4). This finding is in agreement with the concept that RT-DC slightly deforms the cell globally, which means that the main component deformed is the cellular cortex, consisting mainly of actin. The cell area remained smaller in the vimentin-deficient cells, indicating that this parameter is not affected by actin but is determined by vimentin (Fig. 4D).

**Figure 4.**
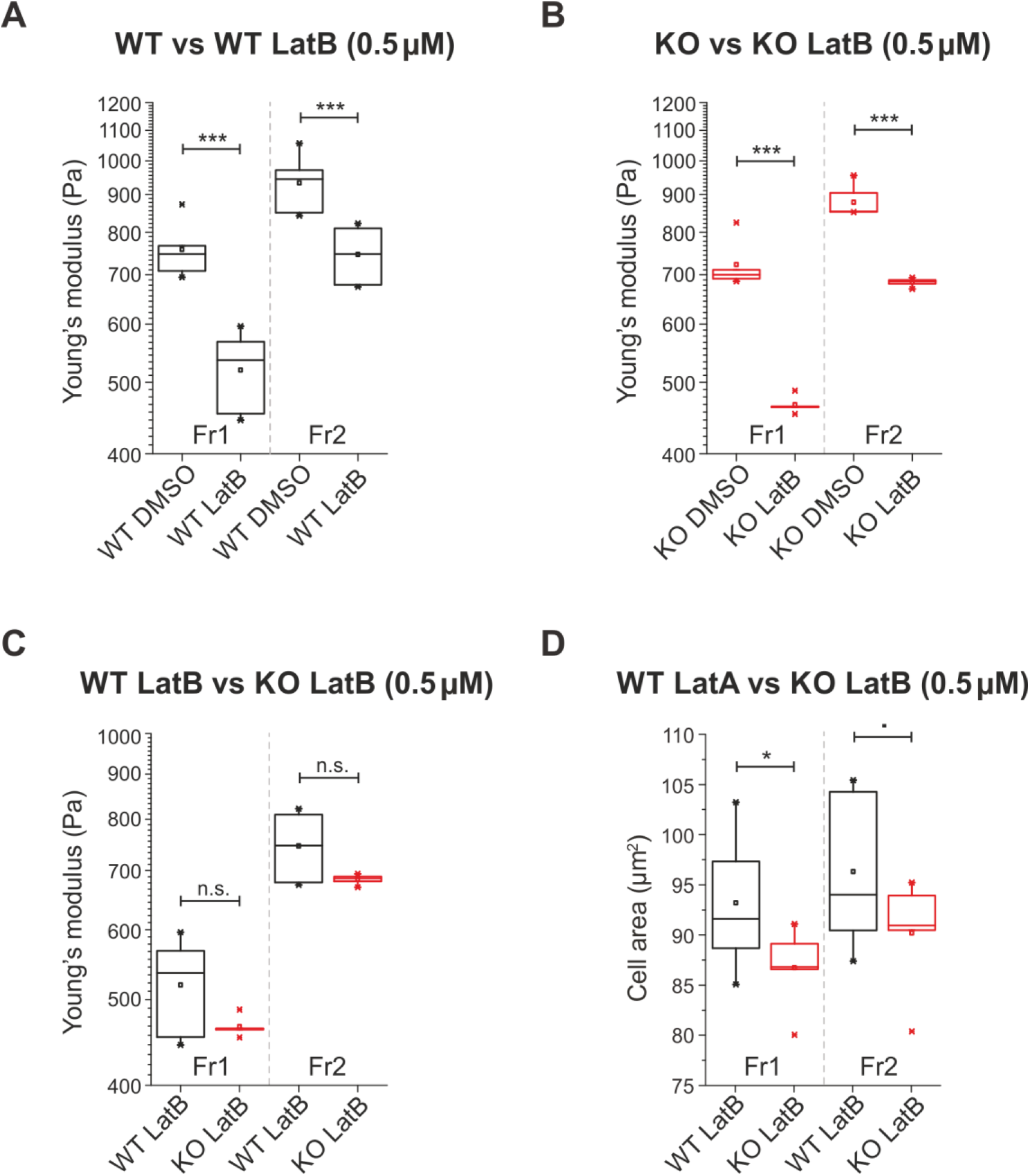
Disruption of microfilaments by latrunculin B decreases cell stiffness on short time scales in both wt and KO cells, as detected by RT –DC measurements. (A-B) Effect on Young’s modulus of 0.5 μM LatB treatment on wt (A) and KO (B) cells. As a control mock DMSO treatment is presented. (C-D) Comparison of Young’s modulus (C) and cell size expressed as cell area (D) of LatB-treated wt and KO cells. Fr1 = 0.16 μl s-1, and Fr2 = 0.32 μl s-1. Box plots of medians from 4 (KO DMSO Fr2) or 5 (rest) independent experiments replicates are shown, boxes present 25th to 75th percentile range, with a line at the median. Whiskers indicate extreme data points within 1.5 × the interquartile range (IQR) . \*\*\**p* < 0.001; * *p* < 0.05, *p*< 0.1; ns, not significant. Statistical analysis was performed using a linear mixed effects model.

We then used AFM, which allows stronger deformation and, therefore, deforms more than only the outer layer of the cells and enabling us to measures the overall mechanical properties of the cells. In contrast to RT-DC data, AFM data showed that only normal control cells, but not vimentin-deficient cells, have a significantly reduced Young’s modulus after the loss of filamentous actin (Fig. 5 A-C, table S5). Vimentin-deficient cells showed no differences in the elastic modulus after the depolymerization of actin for the first two indentations and even an increase in the Young’s modulus for larger indentations resulting in a more prominent deformation of the cells (Fig. 5B). This is contrary to the AFM deformation of cells where actin is intact (Fig. 3), where vimentin-deficient cells were consistently less elastic than control cells. We did not detect any differences in the viscous properties of vimentin-deficient cells upon loss of filamentous vimentin, as compared to control cells (Fig. 5D), which is in agreement with the results described above.

**Figure 5.**
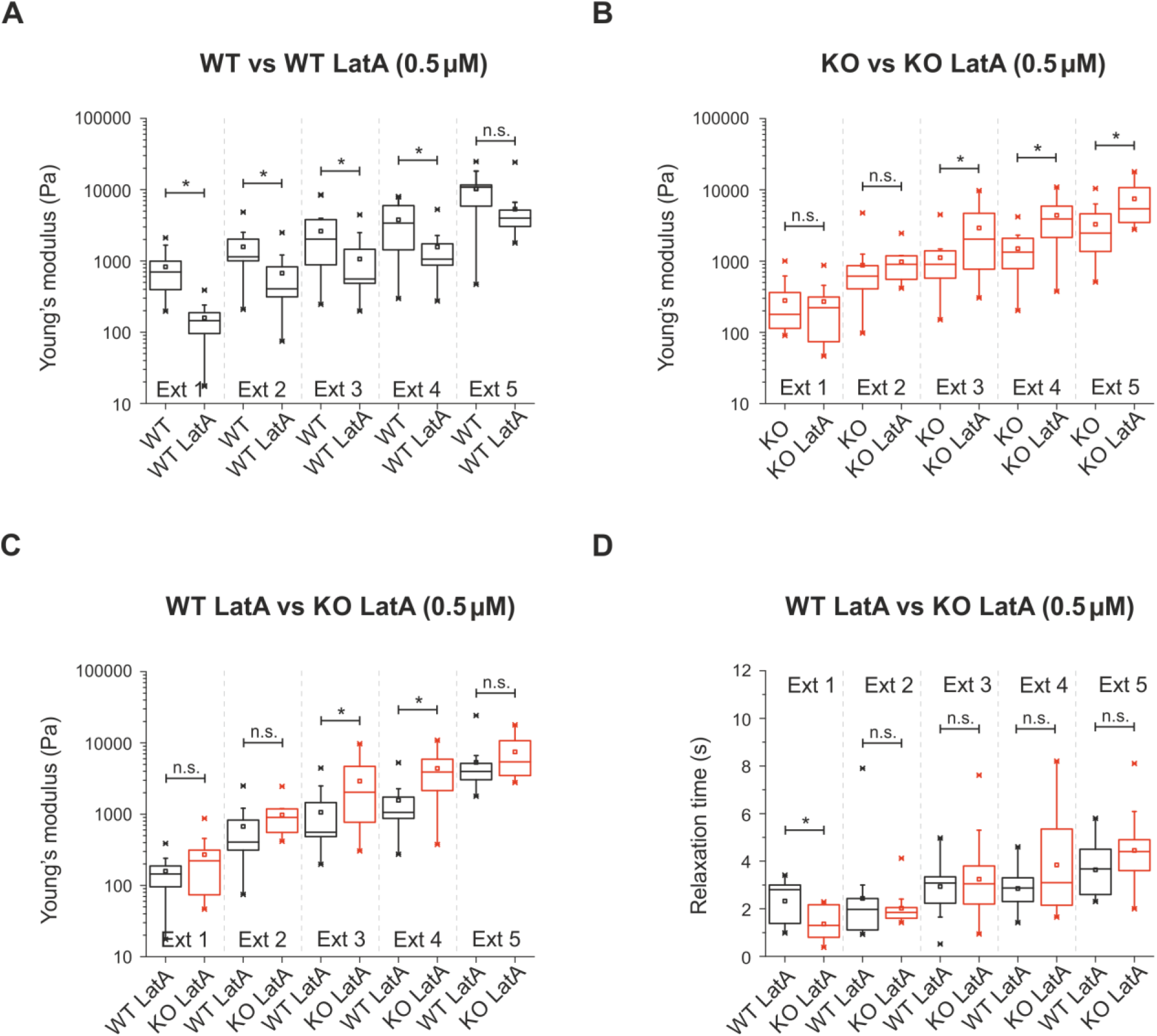
Latrunculin A-mediated microfilament depolymerization disrupts cellular stiffness on long time scales of wt and KO cells differently depending on the presence of vimentin, detected by AFM. (A) Young’s modulus of BMDC wt treated with Latruculin A measured at different extends. (B) Young’s modulus of BMDC vimentin KO treated with latrunculin A measured at different extends. (C) Direct comparison of Young’s modulus of wt (black) and vimentin KO (red) BMDCs treated with Latruculin A measured at different extends. (C) Relaxation time of wt and vimentin KO BMDCs measured at different extends. Results of 3 independent experiments. * *p* ≤0,05. Whisker middle line represent median values. In boxplot mean values are represented by a square.

Importantly, these results demonstrate that it is not the vimentin IFs *per se* that provide the required stiffness at the perinuclear area for dendritic cells but that an interaction between vimentin IFs and filamentous actin provides the required stiffness.

### Vimentin promotes cortical F-actin recovery

The observations above demonstrate that there is an interplay between filamentous actin and vimentin, which plays a role in the control of the mechanical properties of the cell cortex of BMDCs and, therefore, in their migration capacity. In order to further understand the interplay of vimentin with the cell cortex, we analyzed the recovery of cortical actin in vimentin-silenced and control cells, using fluorescence recovery after photobleaching (FRAP) as a method to analyze the subunit exchange of microfilaments. FRAP analysis of microfilament requires live cell imaging of exogenously expressed monomeric actin tagged with fluorescent proteins, which is hard to achieve in primary BMDCs, as they are difficult to transfect. Therefore, we instead transfected hTERT-RPE1 cells with GFP-tagged beta-actin and performed the FRAP analysis of actin in the cortex of the suspended RPE1with normal and reduced amounts of vimentin (Fig. 6 A-C, table S6). By fitting a second order exponential function as suggested by [25, 26] to the recovery curves, we identified two actin populations within the cortex; a slow and a fast recovering population with respective mobile fractions. It has been previously suggested that the slowly and fast recovering population are dependent on formins or Arp2/3, respectively [25, 26]. We then compared the halftime recovery of F-actin in vimentin-silenced cells and control. We observed that reduced vimentin resulted in slower recovery time in both the fast and the slow population of actin (Fig. 6D, E). This decreased capacity to recover cortical F-actin after photobleaching implies that the presence of vimentin promotes the actin turnover at the cellular cortex. The FRAP data also revealed that the mobile fraction of actin monomers was increased in the rapid actin population (Fig. 6 F), whereas it was decreased in the slow actin population in vimentin-reduced cells, as compared to control cells (Fig 6G)–indicating a reduced incorporation of exchangeable actin subunits into actin polymers.

**Figure 6.**
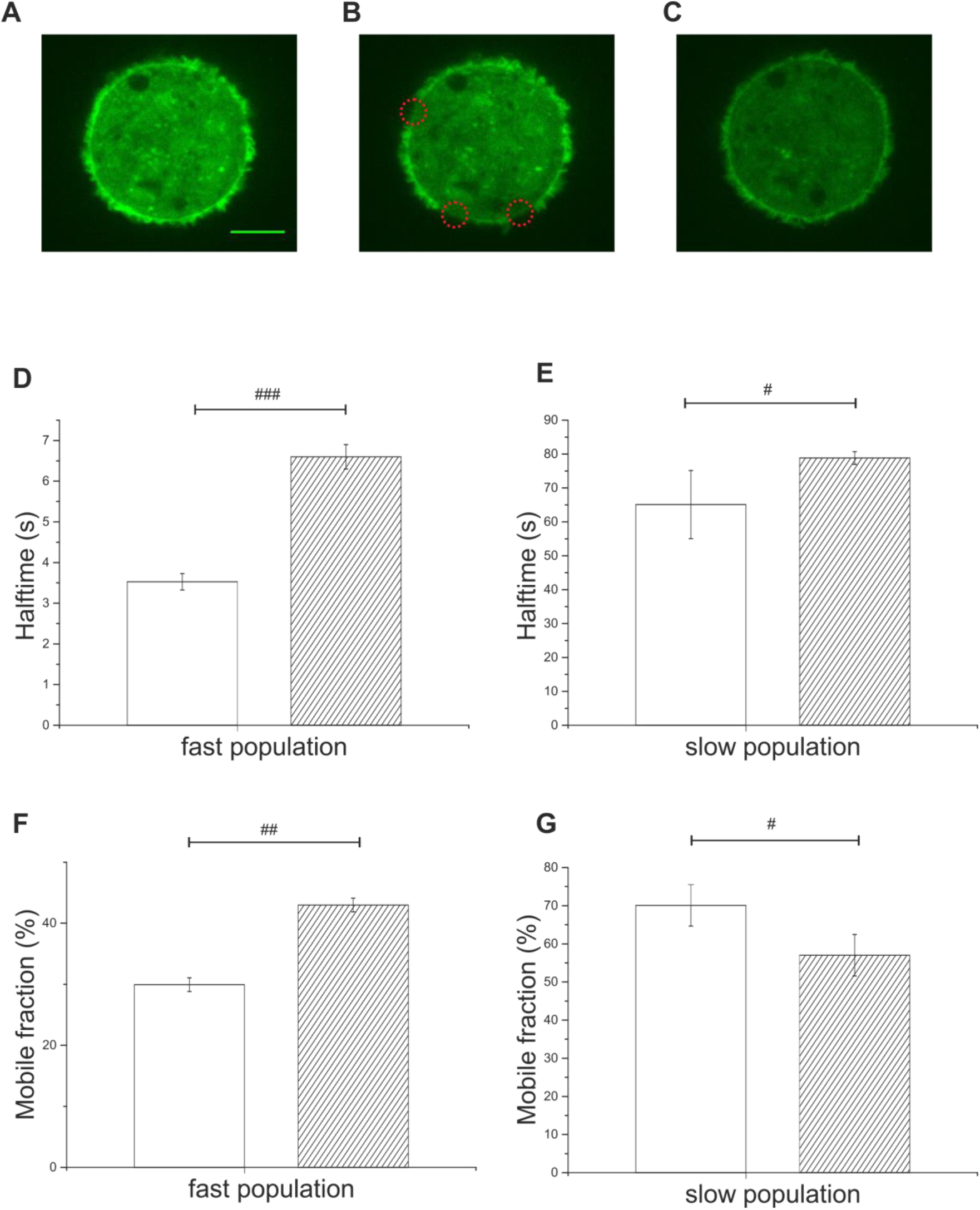
Vimentin promotes cortical F-actin recovery. Representative example of a suspended RPE1 cell (A) before bleaching, (B) directly after bleaching and (C) 80 s after bleaching. (D) Halftime recovery reveals two separated F-actin populations, a slowly (D) and a fast (C) recovering population in vimentin-silenced RPE1 cells (patterned boxplots) and controls (nonpatterned boxplots). The recovery rate of vimentin silenced cells is about 0,5 and 0,2 times slower than of control cells both in the fast and slow population. Percentage of F-actin mobile fraction of the fast (F) and slow (G) population in control (nonpatterned boxplots) and vimentin silenced cells (patterned boxplots). 13 ROIs in 7 WT cells and 10 ROIs in 5 vimentin-silenced cells were measured and analyzed. Data were analyzed for the magnitude of the effect using the Hedges’g test. ### equals large effect d > 1.1, ## equals medium effect d > 0.5, # equals small effect d > 0.2.

All in all, the results demonstrate that vimentin promotes actin dynamics and actin subunit exchange, which is in line with the proposed function of vimentin as an important mediator of migratory capacity.

### Vimentin protects the nucleus from DNA double-strand breaks

We hypothesized that the observed softening, the lower *in vitro* migration and the reduced capacity to reach the lymph nodes *in vivo* in vimentin-deficient BMDCs was due to a loss of mechanical integrity in these cells. There are a number of observations indicating that nuclear positioning is important for migration through confined spaces and that the direction of amoeboid migration governs the positioning of the nucleus [27]. Furthermore, the nuclear positioning is also important for protecting the nucleus from damage that may occur as a result of deformation during the inevitable squeezing that occurs in this type of migration [28]. Hence, the concerted action of vimentin IFs and microfilaments could also facilitate both nuclear positioning and nuclear protection. In this regard, our observation that vimentin mainly has a perinuclear localization (Fig. S1) and also exhibits it mechanical effects primarily at the central, perinuclear region of cells, made us hypothesize that vimentin protects the nucleus and the genome upon mechanical stress. To test this hypothesis, we confined both vimentin-deficient cells and control cells for 24 h in our 2D roof migration setup and analyzed subsequently the DNA double strand breaks of the genome. The data showed that the level of DNA double strand breaks was significantly higher in vimentin-deficient cells, as compared to control (Fig. 7, table S7). In contrast, a DNA double strand break analysis of non-confined cells outside of the confining area in the dish showed no difference in DNA double strand breaks between vimentin deficient cells and control (Fig. 7). We then analyzed the migration of vimentin-deficient and control cells using 1D channels with narrow constrictions, in which the cells were forced to squeeze and deform their full body including their nucleus to pass (Fig. 8A). In addition to the observations related to nuclear stability, we observed that vimentin-deficient cells were slightly blocked or delayed at the constrictions (Fig. 8B). They also displayed a reduced disposition to change their direction at the encounter of a constriction (Fig. 8C), as well as an increased migration speed, as compared to control cells (Fig. 8D). These trends, although not statistically significant, are nonetheless perfectly consistent with our observations that vimentin-deficiency results in cells with lower Young’s modulus, as detected by RT-DC and AFM (Fig. 2E, Fig. 3D).

**Figure 7.**
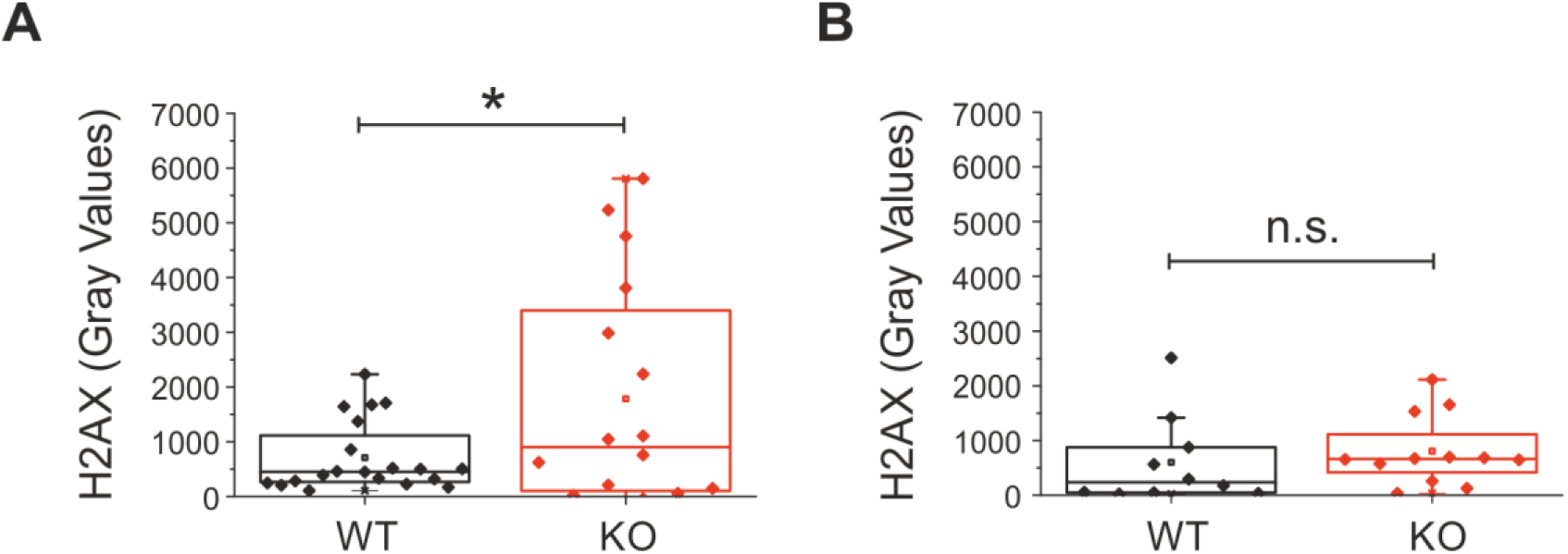
The vimentin network protects the nucleus. DNA double strand break marker H2AX protein levels in the nucleus was measured (A) 24 h after cell confinement under the roof and (B) in the region outside the roof. H2AX levels was measured by immunofluorescence. H2AX intensity in the nucleus were subtracted (normalized) by the intensity obtained in the cytosplasm. Values are displayed in gray values. * *p* ≤0,05.

**Figure 8.**
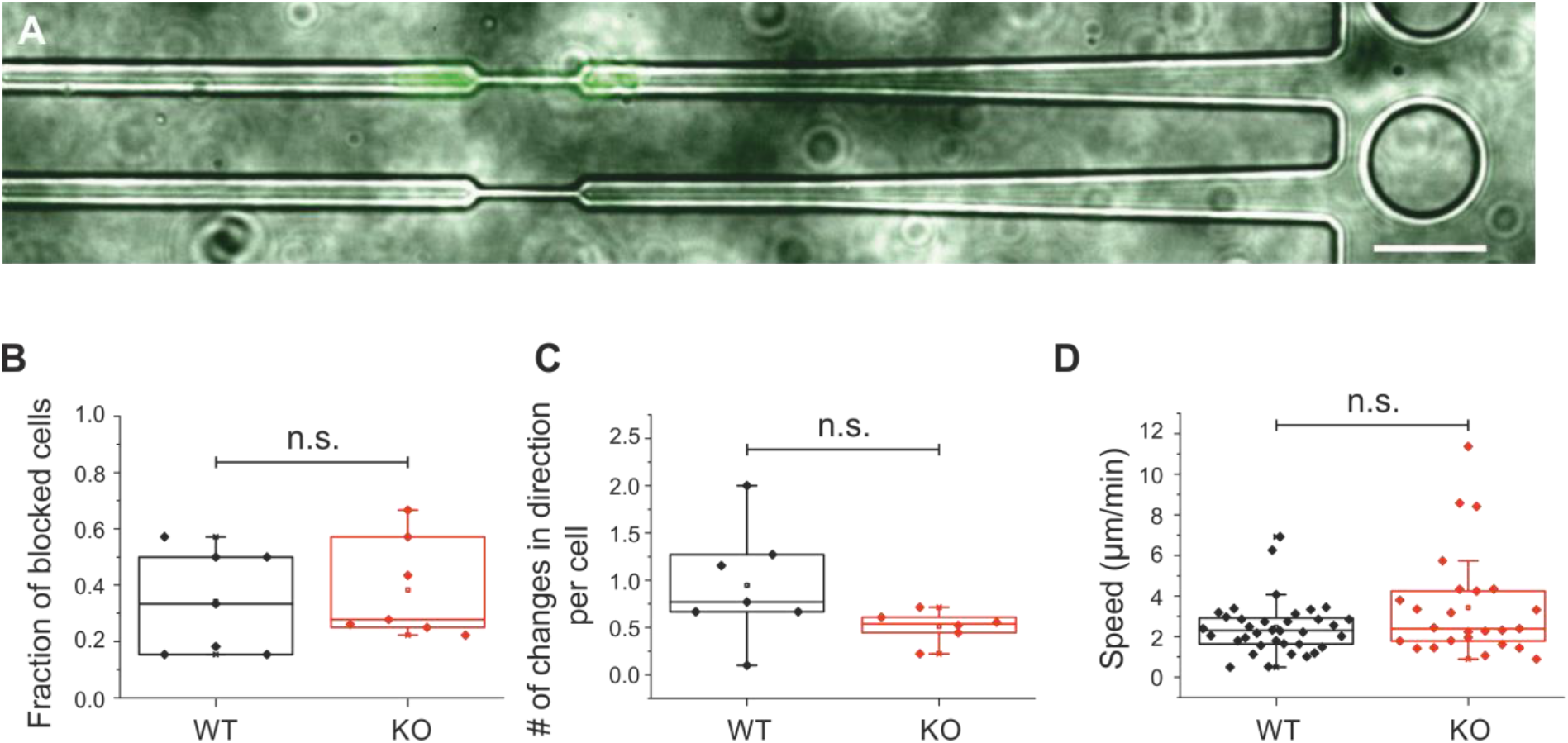
Migration of BMDCs in constricted channels. (A) Immunofluorescence image of constriction channel (phase contrast) with fluorescent image of the nucleus of a BMDC. Scale bar represents 15 μm (B) Ratio between blocked wt (black) and KO (red) BMDCs. (C) ratio of directional changes of wt and KO BMDCs. Each point in the distribution in (B) and (C) represents the ratio between blocked cells /total cells observed per imaged frame during the whole duration of movie (2h). (D) Velocity of cell nucleus migrating through the constriction, each point represents one cell. The total number of cells measured were 36 for wt and 25 for KO cells.

Taken together, these data suggest that vimentin protects cells from nuclear damage when subjected to strong compression. In addition, the proper nuclear positioning may add to the migratory capacity to cells when traveling through confined spaces.

## Discussion

The finding that vimentin controls the motile capacity of cells is in accordance both with the concept that cell migration is governed by the mechanical properties of cells [20] and with the finding that loss of vimentin impairs immune cell migration [9]. It is important to note that the migratory defects observed in vimentin-deficient cells in our *in vitro* experiments were reflected as a major reduction in the capacity of dendritic cells to reach the lymph nodes *in vivo*. This can be explained by the fact that immune cells have to frequently squeeze through numerous tight spaces that they encounter on their way to the lymph node. In this way, the in vitro effect will be both magnified and multiplied.

Confined cells are subjected to nuclear damage [29], and our findings suggest that vimentin protected the DNA from double strand breakage under cellular confinement. Related to protection of DNA against mechanical damage, it has been shown that nuclear positioning is also important for protecting the nucleus from deformation damage [28]. Moreover, the direction of amoeboid migration governs the positioning of the nucleus that in turn directs the amoeboid cells along the path of lowest resistance [27]. As migratory functions and nuclear positioning are intimately coupled, the observed nuclear damage is likely to reflect uncoupling of both nuclear positioning and protection from the migration-induced deformation. This protective effect is expected to manifest itself especially on long time scales, in the order of days but the protection does not necessarily play a major role during circumstances involving rapid and transient migration. This finding is consistent with *in vitro* studies suggesting that the mechanical properties of purified vimentin under compression can have protective effects [30].

In order to investigate the role of vimentin in the migration of immune cells, we studied confined, amoeboid migration using both short- and long-term migration techniques, the latter compatible for more than 2 days. In order to confine the cells in a reproducible manner and to acquire large data sets of trajectories, we used well established protocols for migration in 1D and 2D using microfabrication methods [22, 31]. This allowed us to place our migration data in the landscape of existing amoeboid migration data, e.g. our measured cell speed of WT BMDCs in microchannels of 5.5 μm/min is well comparable to the reported 5 μm/min for the same cell type and treatment [15]. Migration in 2D added further parameter to study such as the migration path length and the apparent persistence, both of which are influenced by the microchannel dimensions in the 1D experiments.

Our observation that loss of vimentin in suspended cells results in softer cells is consistent with previous observations that vimentin is required for the mechanical properties that have been previously characterized only in adherent cells (as reviewed in our review [1]). Related to this, our findings clearly show that vimentin supports cellular elasticity and protects against mechanical stress also in suspended cells, including the compression and squeezing that confined cells are subjected to. Effective migration in confined environments requires surface alterations that involve dynamically orchestrated stiffening of the regions that are responsible for generating the friction forces with the environment that are responsible for pushing the cell forward in the confined space, as it is proposed in [7]. Cells that are pliable, spongy, and yielding, as vimentin-deficient cells will be, are not likely to be able to efficiently generate such forces. This outset for suspension cells is compatible with previous reports regarding adhesive cells, concluding that vimentin IFs serve as a load-bearing scaffold to shield traction stress during single-cell migration [4]. In this study, in the presence of vimentin, actomyosin-dependent traction forces were redirected to peripheral adhesions, which also fits with the concept we are presenting for suspension cells. Our results on actin dynamics also align perfectly with the suggested function of filamentous vimentin in restraining retrograde actin flow [2].

Our finding that loss of F-actin in vimentin-deficient cells did not change the mechanical stiffness of the perinuclear area of cells, indicates that it is not the vimentin IFs or actin filaments *per se* that provide the mechanical cues for perinuclear stiffness, but rather that the perinuclear stiffness depends on an interaction between vimentin and actin that jointly provides the stiffness required for cell migration.

Taken together, our data demonstrate that vimentin collaborates with actin to provide both the elasticity and stiffness needed for amoeboid migration and the perinuclear stiffness required to protect and position the cell nucleus from damage when subjected to the strong compression of amoeboid migration . All of these effects will facilitate the motile capacity of migrating dendritic cells.

## Materials and Methods

### Cells

Primary Dendritic cells were differentiated from bone marrow precursors extracted from vimentin WT and KO mice. BMDCs were generated as previously described in [32] in IMDM medium containing FCS (10 %), glutamine (2 mM), penicillin streptomycin (100 U ml^−1^), 2-ME (50 μM) and further supplemented with granulocyte-macrophage colony-stimulating factor (50 ng ml^−1^) - containing supernatant obtained from transfected J558 cells. The semi-adherent fraction, corresponding to the CD86^+^ cells were gently flushed from the cell culture dishes. All the experiments were performed during cell differentiation day 10-12. All reagents used for cell culture were from Thermo Fisher (Waltham, MA, USA).

Immortalized retinal pigmented epithelium (hTERT-RPE1) cells were cultured in DMEM/F12 medium (Gibco) with 10% fetal bovine serum (Fisher Scientific), 1% GlutaMAX (Fisher Scientific) and 1% penycilin/streptomycin (Gibco).

### Mice

Vimentin heterozygous mice (129/Sv × C57BL/6) were used to generate vimentin-deficient homozygotes (VIM^−/−^) and WT offspring. Both vimentin wt and KO mice used for bone marrow extraction were maintained at the animal facility of the Biocity, Turku, Finland under permit *7284/04.10.03/2012* the protocol number *197/04.10.07/2013* of the Ethical Committee for Animal Experiments of the University of Turku. The regular molecular cloning and other GMO1 related research work was performed under the permit of the Board of Gene Technology, Finland (GTLK) with notification number *038/M/2007*.

### Antibodies and Chemicals

The antibodies used in this study were vimentin monoclonal antibody (D21H3) (Cell Signalling Technology, Danvers, MA, USA), phospho Histone H2A.X (D17A3) (Cell Signalling Technology, Danvers, MA, USA) and CD86 B7-2 (GL-1) (Santa Cruz Technologies) combined with Alexa 647 and Alexa 488 conjugated secondary antibodies (Abcam, Cambridge, UK). For Actin staining we used phalloidin conjugated with tetramethylrhodamine B isothiocyanate (Sigma Aldrich, St Louis, MO, USA) and for nucleus staining Hoechst 34580 (Sigma Aldrich, St Louis, MO, USA). Actin depolymerizing drug latrunculin A (Sigma Aldrich, St Louis, MO, USA) was used at the concentration 0.5 μM Lat A and when applied incubated with the cells throughout the whole experiment. Latrunculin B was used in RT-DC measurements.

### Channels

Microchannels used in migration experiments were manufactured according to previously published procedures [22, 33], and for the experiments using channels with constrictions we used the design described by [34]. Briefly, silicone rubber RTV-615 kit (Momentive performance materials, US) was mixed in a proportion 10:1-silicone/curing agent, degassed and polymerized at 75°C in the specific microfabricated molds containing the channel positive imprint. The resultant channels of 5 μm height and 5 μm width was attached in 35mm glass bottom cell culture dishes (World Precision Instruments, Sarasota, FL, USA) by plasma surface activation. The assembled structure was coated with Fibronectin (200 mg mL^−1^) (Sigma-Aldrich, St Louis, MO, USA) and incubated with cell culture medium for at least 30 min. Cells were platted in the channels entry at a concentration of 2×10^7^ cells mL^−1^.

### Constriction channels

Constriction channels were produced as straight channels using a different template. The constrictive part was 15 μm long and 2 μm wide, see [34] for details. We calculated the ratio of cells being blocked at the constriction, how many cells changed their direction at the encounter of the constriction as well as the speed of the nucleus when passing the constrictions. Each point in Fig 6B, C represents the ratio between blocked cells and the total cells observed per imaged frame during the whole duration of one movie (2h) with one image taken every 2 min. In Fig 6d each point represents one cell.

### Plate-plate confiner

Confining cell roofs were prepared as previously described [17, 18]. The mold for the coverslip PDMS coating was produced by photolithography and consists of pillars of 3 μm height and 440 μm diameter, spaced 1000 μm from each other. The confiner was assembled in a glass bottom 6 well plate (Mattek, Ashland, MA, USA). The roof of 3 μm height was coated in a 10 mm coverslip. The roof was stitched to the culture dish lid by a softer, deformable PDMS pillar that was later closed and fixed with adhesive tape. Before the experiments, both cell culture dishes and the confiner roof were coated with PLL-PEG (0.5 mg/mL) (SUSOS, Dübendorf, Switzerland) to avoid cell adhesion. The mounted device was incubated prior to the experiment in culture medium for at least 4h to equilibrate the PDMS. Cell recording started 2h after cell confinement.

### Migration assays cell trajectories

Cell nucleus were stained with Hoechst 34580 (200 ng/mL for 30 min) (Sigma Aldrich, St Louis, USA) and migration was recorded by epifluorescence microscopy for at least 6h. Cells were kept in a constant atmosphere 37°C and 5%CO2 during the entire experiment. Images were obtained using the inverted microscope system eclipse Ti-E (Nikon, Tokyo, Japan) equipped with a fluorescent illumination. Cell trajectories were tracked using the custom-made software described in [35, 36]. For analysis, all trajectories were filtered to exclude trajectories shorter than 50 μm or 30min.

### Apparent persistence

The apparent persistence was calculated dividing the diameter of a circle including the full trajectory by the full length of the trajectory. This value would be 1 if the cell migrates perfectly straight and <1 for less persistent cell trajectories.

### *In vivo* lymph-node homing assay

*In vivo* migration assay was performed according to [37] with smaller modifications. Prior to mice injection, bone marrow differentiated dendritic cells from vimentin KO and wt mice were treated by LPS (100 ng/mL for 30 min) and fluorescently labeled with Cell tracer Oregon 488 (Thermo Fisher Scientific, Waltham, MA, USA), according to the manufacturer’s protocol. A total of 1×10^6^ fluorescently tagged cells, diluted in PBS1X in a final volume of 40 mL were subcutaneously injected into the hind footpad of mice aged between (6-10 weeks old). Mice were killed 36 h after BMDC injection and both popliteal and inguinal lymph nodes were extracted. The lymph nodes were then mechanically disrupted and digested by collagenase D (Roche, Basel, Switzerland) and DNAse1 (Roche, Basel, Switzerland) for 30 min at 37·C. The homogenate was filtered in a 77 μm silk filter. The purified cells were mounted in a microscope slide and the ratio between fluorescent and non-tagged cells was counted. This ratio was normalized to the ratio of WT cells arriving in the lymph node.

### Immunofluorescence

Cells were fixed for 10 min with Paraformaldehyde solution 4% (Sigma Aldrich, St Louis, MO, USA) and permeabilized with Triton 0.5% (Sigma Aldrich, St Louis, MO, USA) for 10 min. Protein blocking was done by incubating cells with Bovine serum albumin (BSA) 3% solution for 1 h. Primary antibodies were diluted at 1:200 ratio in BSA 3% and incubated overnight at 4°C. Secondary antibodies were diluted at 1:1000 ratio in BSA 3% and incubated for 1h at room temperature. Actin staining was done by Phalloidin conjugated with Rhodamine (1mM final concentration).

Cells were mounted using Moewiol (Sigma Aldrich, St Louis, MO, USA) or Fluoromount G containing Dapi (Invitrogen, Carlsbad, CA, USA) to stain the nucleus. For H2AX quantification: The staining intensity of H2AX and DAPI in the nucleus were measured in gray scale and normalized by the values obtained in the cytoplasm.

### SDS-PAGE and western blot

Whole cell lysates were extracted by Laemmli sample buffer. Protein separation was done by SDS-PAGE in 10% bis-acrylamide gels and transferred to the nitrocellulose membrane Amersham Protran, pore size 0.45 μm (GE Healthcare, Chicago IL, USA). Membrane was blocked with 5% fat free milk in TBS and incubate overnight at 4°C with Vimentin antibody (dilution 1:1000 in BSA 5%) or with the loading control HSC70 (dilution 1:1000 in BSA 5%).

### Confocal microscopy

Confocal images were obtained using a Yokogawa spinning disk unit (CSU W1, Andor Technology, Belfast, UK) with a pinhole size of 50 μm coupled to the inverted microscope system Eclipse Ti-E (Nikon, Tokyo, Japan). Images were recorded using a Hamamatsu flash 4.0 camera with a 6.5 μm pixel size (Hamamatsu, Hamamatsu city, Japan). Image treatment and Z maximum projection were done using the software ImageJ FIJI [38].

### Atomic force microscopy

To measure the global stiffness of cells in suspension, tipless cantilevers (MLCT-010, cantilever F) with a nominal spring constant of 0.3-1,2 (nominal: 0,6) N m^−1^. (Bruker, Billerica, MA, US) were wedged to correct the 10° cantilever tilt, resulting in a flat surface ton probe cells. Wedged cantilevers were done according to [24]. Tipless cantilevers were pressed against drops of Norland Optical Adhesive 63 (NOA63) (Thorlabs, Newton, NJ, US) placed on a silicon coated coverslip. NOA63 was cured by UV light for 60 s, gently detached from the silicon coverslip and cured for additional 300 s. Flatness and integrity of wedged cantilevers was accessed by electron microscopy.

Atomic force microscopy (AFM) measurements were done using JPK Cell Hesion 200 (JPK Instruments, Berlin, Germany), mounted on the inverted microscope system eclipse Ti-E (Nikon, Tokyo, Japan). Sensitivity was calibrated by acquiring a force curve on the dish surface and spring constant was calibrated by the thermal noise fluctuation method using JPK build in software. For measurements, cantilever was lowered at speed 0.5 μm/s set point force was set at 4nN. Measurements consisted of 5 subsequent compressions of 1 μm for 60 s each. During compressions, height was kept constant and cell force was recorded for all extends (total time: 300s). Bright-field image were taken in all steps of the measurement (magnify 40X plus 1,5 zoom in order to calculate the size and contact area of the cells absent in cellular protrusions). Cells were kept in at 37°C during measurements and culture media was supplemented with Hepes buffer 200 mg/mL. The Young’s modulus was calculated using the Hertz-approximation on each extend [39]. The Hertz-model is a purely elastic model, which means that the here depicted “Young’s modulus” is basically an apparent elastic modulus, because cells are viscoelastic objects.

### Real-time deformability cytometry (RTDC)

High throughput measurements of cell mechanics were performed using RT-DC according to previously described procedures [40]. In brief, cells were suspended in 100 μl of a viscosity-adjusted measurement buffer (0.5% (w/v) methylcellulose (4000 cPs, Alfa Aesar, Germany) in phosphate saline buffer without Mg^2+^ and Ca^2+^; final viscosity 15 mPa s) and flushed through a 300-μm long microfluidic channel with a 30 × 30 μm square cross-section at flow rates of 0.16 μl s^−1^ (Fr1) and 0.32 μl s^−1^ (Fr2). The images of deformed cells were acquired at 2,000 frames s^−1^ within a region of interest close to the channel end. Cell deformation and cross section area were evaluated in real-time based on contours fitted to cells by an image processing algorithm developed in house [23]. Obtained data were filtered for cell area of 50–500 μm^2^ and the area ratio of 1.00–1.05. Area ratio is defined as the ratio between the area enclosed by the convex hull of the contour and the area enclosed by the raw contour, and allows for discarding the cells with rough or incompletely fitted contours. Young’s modulus values were assigned to each cell using a data grid obtained from numerical simulations for an elastic solid [41] with the aid of the analysis software ShapeOut version 0.8.7 (available at https://github.com/ZELLMECHANIK-DRESDEN/ShapeOut). A minimum of 500 cells were analysed per condition in each experiment. Statistical analysis was performed using linear mixed effect model as described in details elsewhere [42].

### Actin dynamics

Actin cortex dynamics was tested by fluorescence recovery after photobleaching (FRAP) on actin cortex as described in [26] (FRAPPA, Andor Technology). We simulated the cortex of BMDC cells using suspended RPE1 cells. In these cells, we fluorescently labeled beta actin monomers using CellLight Reagent BacMam 2.0 GFP (by Life technologies). Additionally, vimentin was transfected using m-Cherry fluorescent plasmids and silenced using 10 μM diluted Vimentin Silencer siRNA (life technologies) and Lipofectamine RNAiMAX Reagent (ratio 1:1), both diluted in serum free medium. Cells were incubated in a normal incubation conditions (37°C, 5% CO2) for 3 days, the silencing process was repeated after 3 or 4 days.

For bleaching, 100% of the maximum laser power (50 mW) were applied to a circular region of interest (ROI) 2μm in diameter on the actin cortex of control and vimentin silenced cells. ROI fluorescent recovery was imaged with 448nm wavelength for 80-100 s after bleaching using a confocal microscope (Nikon). Intensity values of the fluorescence within the selected ROI, the reference ROI (normally in the cytoplasm), and the background ROI were analyzed using the image processing software ImageJ. We used these ROIs to correct the raw data for overall bleaching of the cell. The intensity values of the corrected data were normalized between 0 and1 (1 before and 0 directly after the bleaching pulse). By fitting a second order exponential recovery function as suggested by [25, 26] to the mean recovery curve we identified two actin populations within the cortex — a slowly and a fast recovering population with respective mobile fractions. It has previously been suggested that the slowly recovering population is formin mediated, the fast recovering population Arp2/3 mediated [25, 26].

## Acknowledgements

The Authors would like to thank Kevin Kaub for helpful discussions concerning the FRAP measurements.

