## Supplementary Information for "Vimentin provides the mechanical resilience required for amoeboid migration and protection of the nucleus"

***Supplementary figure + legend***

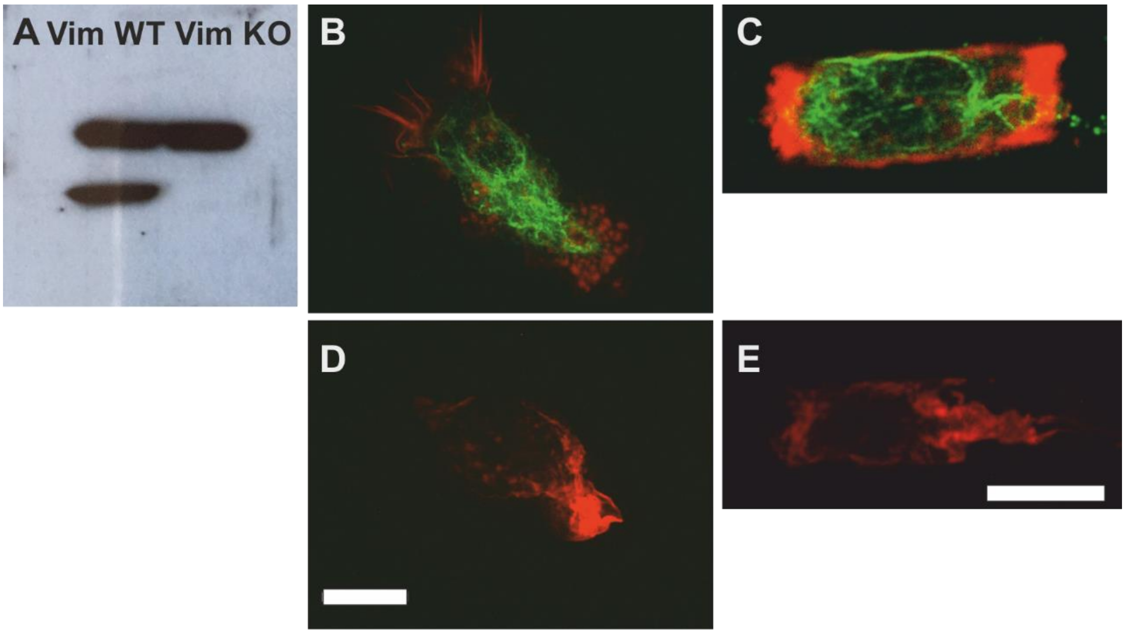


**Fig S1**: (A) Protein quantification of vimentin in wt and vimentin KO primary BMDC by Western blot (vimentin 54 kDa). Loading control HSC70 (70 kDa). B-E; Representative images of actin (red) and vimentin (green) on a glass coverslip and in 5 mm high and 5 mm wide micro-channels with all surfaces coated with fibronectin showing wt (B-C) and (D-E) vimentin KO BMDC. Scale bars represent 10 mm.

*Supplementary tables*

**Table S1. Loss of vimentin results in defective amoeboid migration.**

| System | Observable | Cell type | Mean value ± SE |
|---|---|---|---|
| 1D | Migrating cells | WT | 72.2 ± 2.4 |
| | (%) | KO | 57.8 ± 5.3 |
| | Migration velocity | WT | 5.55 ± 0.09 |
| | (µm/min) | KO | 4.98 ± 0,11 |
| | Apparent persistence | WT | 0.68 ±0.007 |
| | | KO | 0.66 ± 0.01 |
| 2D | Migrating cells | WT | 52.7 ± 9.14 |
| | (%) | KO | 42.0 ± 10.15 |
| | Migration velocity | WT | 5.5 ± 0.05 |
| | (µm/min) | KO | 4.5 ± 0.05 |
| | Path length | WT | 383 ± 23.6 |
| | (µm) | KO | 240 ± 19 |
| | Apparent persistence | WT | 0.51 ± 0.004 |
| | | KO | 0.51 ± 0.004 |
| In vivo | Cells arriving at the lymph node | WT/WT | 0.54 ± 0.23 |
| | (normalized) | KO/WT | 0.55 ± 0.16 |
| | | WT/KO | 1 ± 0.5 |
| | | KO/KO | 0.24 ± 0.07 |

**Table S2. Loss of vimentin in dendritic cells decreases cell stiffness as detected by RT-DC.** Values represent medians ± MAD (median absolute deviation).

| Flow rate | Cell types | No exp | No cells per exp | No cells total | Deformation | Young's modulus (kPa) | Cell area (µm²) |
|---|---|---|---|---|---|---|---|
| 1 | WT | 5 | 3.069; 2.690; 1.371; 1.614; 1.494 | 10.238 | 0.0256 ± 0.0025 | 776 ± 22 | 90.1 ± 3.6 |
| | KO | 5 | 1.929; 1.991; 2.212; 1.836; 1.526 | 9.494 | 0.0271 ± 0.0012 | 706 ± 12 | 82.9 ± 1.2 |
| 2 | WT | 5 | 1.652; 2.550; 1.055; 1.783; 2.053 | 9.093 | 0.0403 ± 0.0046 | 926 ±66 | 93.1 ± 5.7 |
| | KO | 5 | 2.134; 2.218; 2.512; 1.971; 1.786 | 10.621 | 0.0469 ± 0.0008 | 838 ± 16 | 86.2 ± 1.8 |

**Table S3. Force-mode atomic force microscopy analysis of dendritic cell mechanics.** Values represent mean ± StDv.

| Extend | Cell types | No cells | Youngs Modulus (Pa) | No cells | Relaxation time (s) |
|---|---|---|---|---|---|
| 1 | WT | 12 | 826 ± 575 | 8 | 1.7 ± 1.1 |
| | KO | 16 | 280 ± 248 | 13 | 1.9 ± 1.8 |
| 2 | WT | 12 | 1573 ± 1221 | 7 | 2.6 ± 1.2 |
| | KO | 16 | 878 ± 1073 | 14 | 2.4 ± 1.3 |
| 3 | WT | 11 | 2617 ± 2295 | 8 | 3.1 ± 1.6 |
| | KO | 16 | 1118 ± 993 | 17 | 3.4 ± 1.5 |
| 4 | WT | 10 | 3782 ± 2731 | 7 | 5.7 ± 2.3 |
| | KO | 16 | 1491 ± 991 | 17 | 4.9 ± 2.4 |
| 5 | WT | 9 | 10275 ± 7669 | 6 | 4.9 ± 1.7 |
| | KO | 16 | 3288 ±2583 | 17 | 4.8 ± 1.0 |

**Table S4. Latrunculin-treatment decrease cell stiffness on short time scales in both wt and KO cells. as detected by RT –DC measurements**. Values represent medians ± MAD (median absolute deviation)

| Flow rate | Cell types | No exp | No cells per exp | No cells total | Young's modulus (kPa) | Cell area (µm²) |
|---|---|---|---|---|---|---|
| 1 | WT DMSO | 5 | 3.256; 3.044; 2.030; 1.861; 2.297 | 12.488 | 747 ± 38 | 91.6 ± 3.4 |
| | WT LatB | 5 | 1.177; 1.266; 956; 839; 1.285 | 5.523 | 536 ± 60 | 91.6 ± 5.7 |
| | KO DMSO | 5 | 2.001; 1.975; 2.272; 1.668; 2.177 | 10.093 | 700 ± 11 | 87.5 ± 2.5 |
| | KO LatB | 5 | 729; 909; 959; 786; 947 | 4.330 | 464 ± 1 | 86.8 ± 2.3 |
| 2 | WT DMSO | 5 | 3.329; 1.603; 2.277; 2.218; 2.981 | 12.408 | 944 ± 94 | 93.8 ± 5.2 |
| | WT LatB | 5 | 935; 501; 774; 641; 984 | 3.835 | 747 ± 68 | 94.0 ± 6.6 |
| | KO DMSO | 4 | 919; 2.774; 2.155; 2.384 | 8.232 | 853 ± 1 | 87.1 ± 1.1 |
| | KO LatB | 5 | 574; 546; 740; 648; 821 | 3.329 | 686 ± 5 | 91.0 ± 3.0 |

**Table S5. Latrunculin-treatment effects cellular stiffness on long time scales of wt and KO cells differently depending on the presence of vimentin. detected by AFM.**

| Extend | Cell type and treatment | No cells | Youngs Modulus (Pa) | No cells | Relaxation time (s) |
|---|---|---|---|---|---|
| 1 | WT | 12 | 826 ± 575 | 13 | 2.3 ± 0.9 |
| | WT LatA | 17 | 159 ± 101 | | |
| | KO | 16 | 280 ± 248 | 11 | 1.4 ± 0.7 |
| | KO LatA | 10 | 270 ± 249 | | |
| 2 | WT | 12 | 1573 ± 1221 | 13 | 2.4 ± 2.0 |
| | WT LatA | 18 | 671 ± 618 | | |
| | KO | 16 | 878 ± 1073 | 13 | 2.0 ± 0.7 |
| | KO LatA | 11 | 974 ± 569 | | |
| 3 | WT | 11 | 2617 ± 2295 | 14 | 2.9 ± 1.2 |
| | WT LatA | 18 | 1065 ± 1035 | | |
| | KO | 16 | 1118 ± 993 | 13 | 3.2 ± 1.9 |
| | KO LatA | 11 | 2912 ± 2737 | | |
| 4 | WT | 10 | 3782 ± 2731 | 12 | 2.9 ± 0.9 |
| | WT LatA | 17 | 1566 ± 1442 | | |
| | KO | 16 | 1491 ± 991 | 12 | 3.8 ± 2.1 |
| | KO LatA | 9 | 4389 ± 3431 | | |
| 5 | WT | 9 | 10275 ± 7669 | 13 | 3.6 ± 1.1 |
| | WT LatA | 16 | 5321 ± 5239 | | |
| | KO | 16 | 3288 ± 2583 | 11 | 4.5 ± 1.7 |
| | KO LatA | 8 | 7490 ± 5260 | | |

**Table S6. Fluorescent recovery after photobleaching data for F-actin recovery speed**. Values represent mean ± SE.

| Cell type | Population | No of analysed cells | No of analysed ROIs | Half time (s) | Mobile fraction (%) |
|---|---|---|---|---|---|
| RPE1 | | | | | |
| | Fast | 7 | 13 | 3.53 ± 0.2 | 30.9 ± 1.13 |
| | Slow | 7 | 13 | 65,1 ± 10 | 70,06 ± 5.4 |
| RPE1 VIM - Si | | | | | |
| | Fast | 5 | 10 | 6.6 ± 0.3 | 42.93 ± 1.13 |
| | Slow | 5 | 10 | 78.9 ± 1.8 | 57 ± 5.43 |

**Table S7. The vimentin network protects the nucleus.** Values represent mean ± StDv.

| Location of measurement | Cell type | No | Gray values, Mean ± Stdev |
|---|---|---|---|
| Within confined area | WT | 20 | 709 ± 640 |
| | KO | 16 | 1787 ± 2071 |
| Outside confined area | WT | 10 | 601 ± 811 |
| | KO | 12 | 805 ± 636 |

**Table S8. Migration of BMDCs in constrion channels**. Values represent mean ± StDv.

| Cell type | No | Ratio of blocked cells | No of directional changes/ No of observed cells | Velocity (µm/min) |
|---|---|---|---|---|
| WT | 36 | 0.34 ± 0.18 | 0.30 ± 0.92 | 2.5 ± 1.3 |
| KO | 25 | 0.38 ± 0.17 | 0.26 ± 0.56 | 3.4 ± 206 |
